## Supplemental Fig.1-5 for "Human RAD51 paralogue, SWSAP1, fosters RAD51 filament by regulating the anti-recombinase, FIGNL1 AAA+ ATPase"

### Supplementary Figures

#### Supplementary Fig. 1

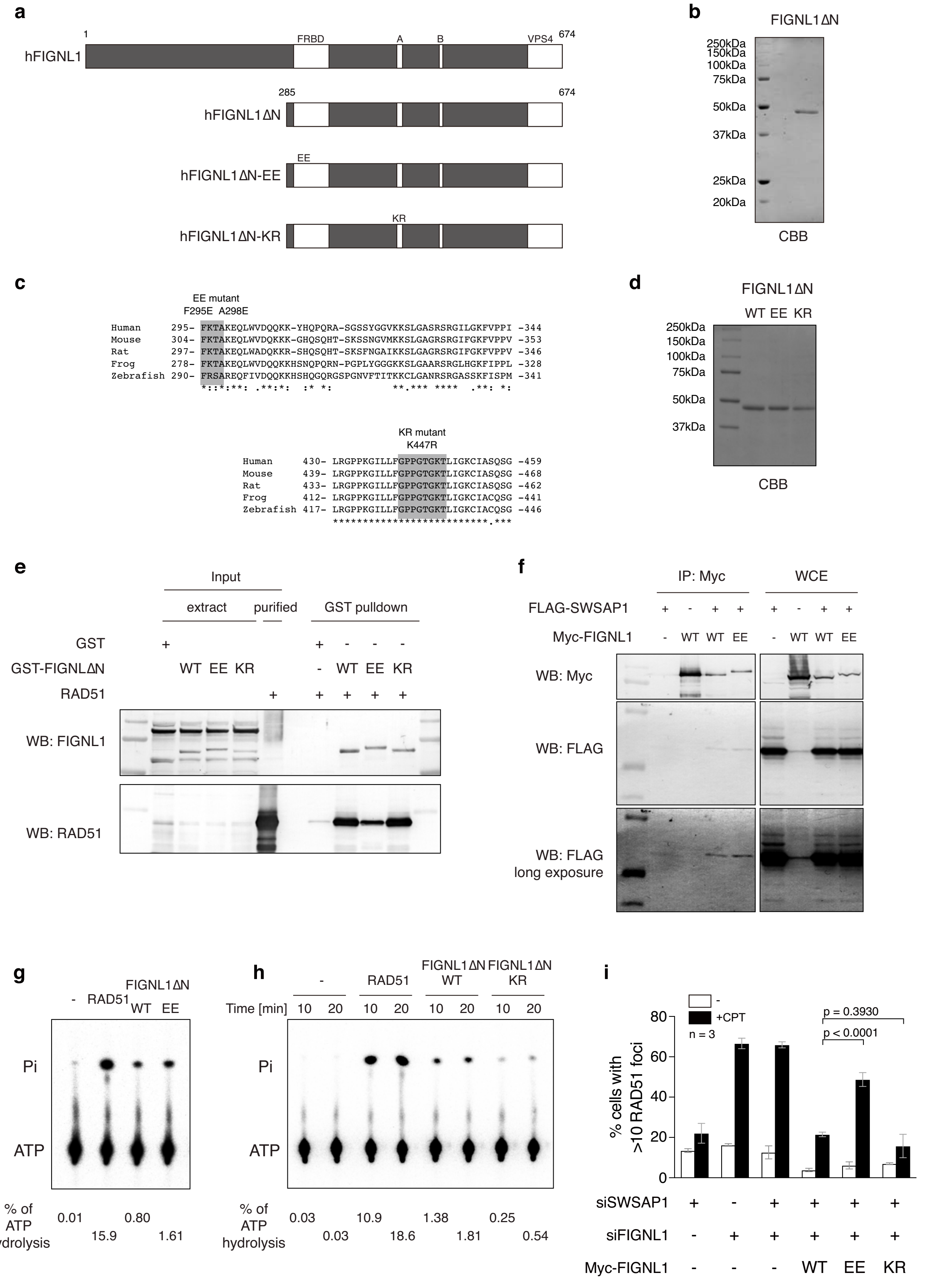


**Supplementary Fig. 1: Functional analysis of FIGNL1 mutants**

**a,** Schematic representation of human FIGNL1, FIGNL1ΔN, FIGNL1ΔN-EE and FIGNL1ΔN-KR used in Fig. 4. **b,** Purified FIGNL1ΔN was analyzed by SDS-PAGE with Coomassie staining. **c,** Amino acid sequence comparison of FIGNL1's RAD51 binding domain (FRBD) and Walker A motif (shaded region). **d,** Purified FIGNL1ΔN-EE and -KR were analyzed by SDS-PAGE with Coomassie staining. **e,** Interaction between purified FIGNL1 mutant proteins and RAD51. Bacterial extract expressing GST, GST-FIGNL1ΔN, GST-FIGNL1ΔN-EE or GST-FIGNL1ΔN-KR were incubated with glutathione beads. After wash, the beads were incubated with purified RAD51. Samples were eluted with glutathione and subjected to western blotting. **f,** Co-immunoprecipitation analysis of FIGNL1-EE mutant. Myc-FIGNL1 or Myc-FIGNL1-EE were co-expressed with FLAG-SWSAP1 in 293T cells, and subjected to IP and analyzed by western blotting with indicated antibodies. **g-h,** ATPase activity of FIGNL1ΔN, FIGNL1ΔN-EE and FIGNL1ΔN-KR was analyzed. [$\gamma$-$^{32}$P]ATP was incubated with indicated proteins. The products were analyzed by thin-layer chromatography using PEI plates. The plates were analyzed by the phosphor imager and its images are shown. **i,** After 96-h transfection of siRNA for FIGNL1 and SWSAP1 with the expression of siRNA-resistant FINGL1, FIGNL1-EE or FIGNL1-KR mutnat proteins, cells were treated with 100 nM of CPT for 22 h and immuno-stained for RAD51. Quantification of RAD51-positive cells was analyzed. Data are mean ± s.d.; Statistics and reproducibility, see accompanying Source data.

**Fig. S2**

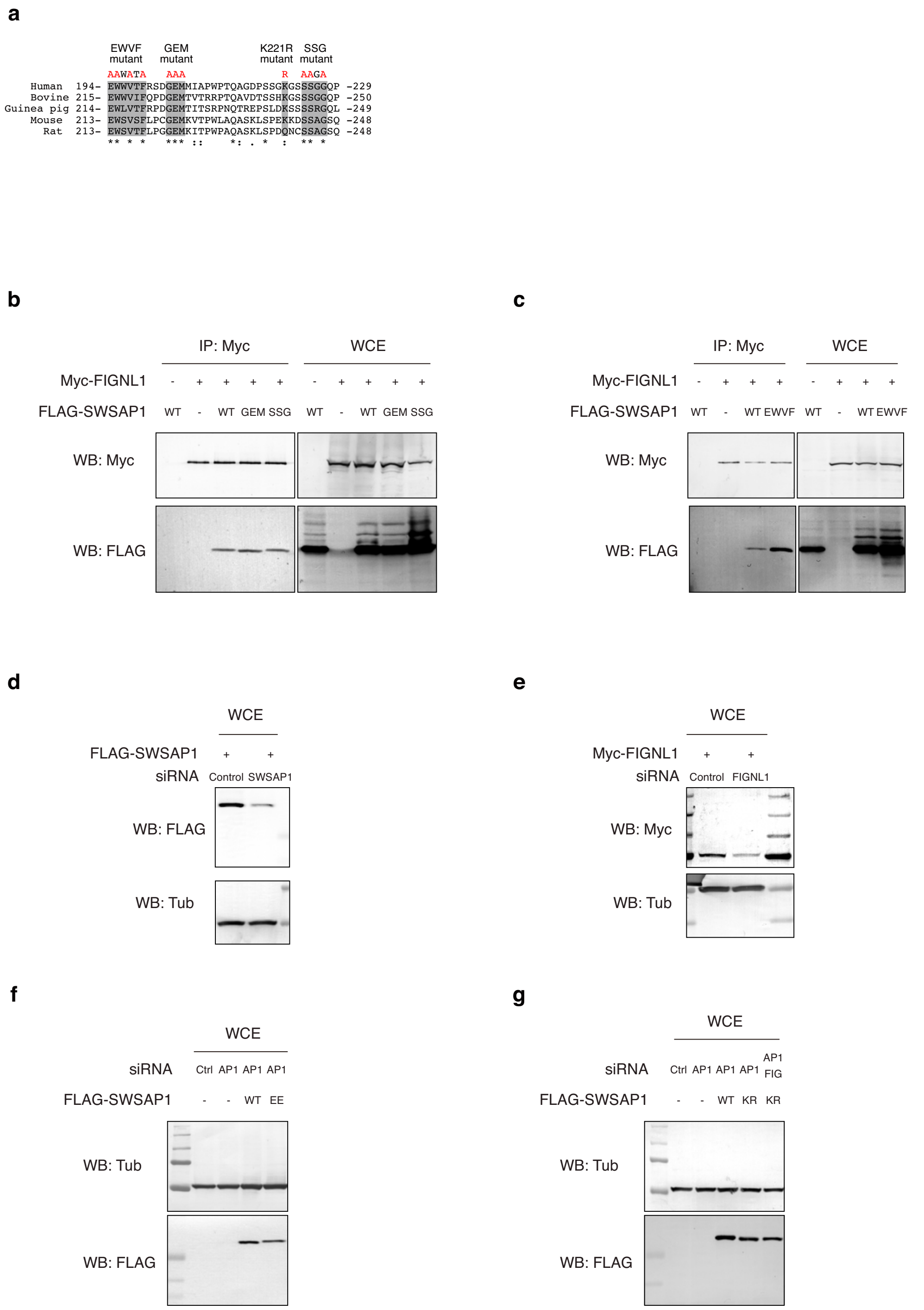


**Supplementary Fig. 2: Interaction between SWSAP1 C-terminal point mutants and FIGNL1**

**a,** Amino acid sequence comparison of conserved SWSAP1 C-terminus. Amino acid substitutions (red) are indicated above the sequences. **b-c,** Co-immunoprecipitation analysis of SWSAP1 mutants. Myc-FIGNL1 and indicated FLAG-SWSAP1 mutant proteins expressed in 293T cells were used for IP and analyzed by western blotting with FLAG and Myc antibodies. **d,** SWSAP1 depletion in FLAG-SWSAP1-expressing cells. Evaluation of the effect on siRNA for SWSAP1 in U2OS cells was analyzed for FLAG-SWSAP1. Tubulin was an internal control. Tubulin was an internal control. **e,** FIGNL1 depletion in Myc-FIGNL1-expressing cells. Evaluation of the effect on FIGNL1 siRNA in U2OS cells was analyzed for Myc-FIGNL1. Tubulin was an internal control. **f,** Protein expression of siRNA-resistant SWSAP1 and SWSAP1-EE. Evaluation of the effect on SWSAP1 siRNA on RNAi-resistant for SWSAP1 and SWSAP1-EE in U2OS cells was examined. Tubulin was an internal control. **g,** Protein expression of siRNA-resistant SWSAP1 and SWSAP1-KR. Evaluation of the effect on SWSAP1 siRNA on RNAi-resistant for SWSAP1 and SWSAP1-KR in U2OS cells was studied.

**Fig. S3**

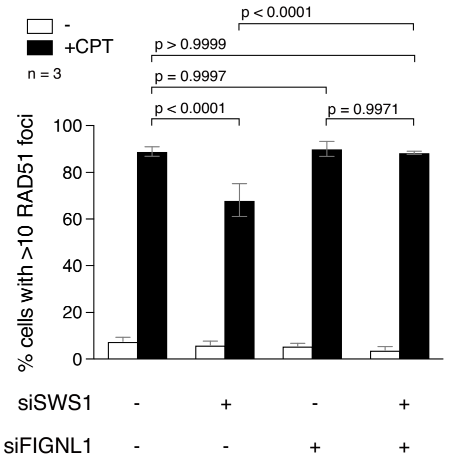


**Supplementary Fig. 3: RAD51 focus formation defect in SWS1-depleted cells is suppressed by the FIGNL1 depletion**

Immuno-staining analysis of RAD51 focus in control U2OS cells and SWS1-depleted cells with or without the FINGL1 depletion (top two raw) at 22 h after the treatment of 100 nM of camptothecin (CPT). Quantification of RAD51 focus-positive cells (more than 10 foci per a cell) in the indicated cell lines. At each point, more than 200 cells were counted. Quantification of RAD51 focus-positive cells in the indicated siRNA-transfected cells. Data are mean ± s.d.; Statistics and reproducibility, see accompanying Source data.

**Supplementary Fig. 4**

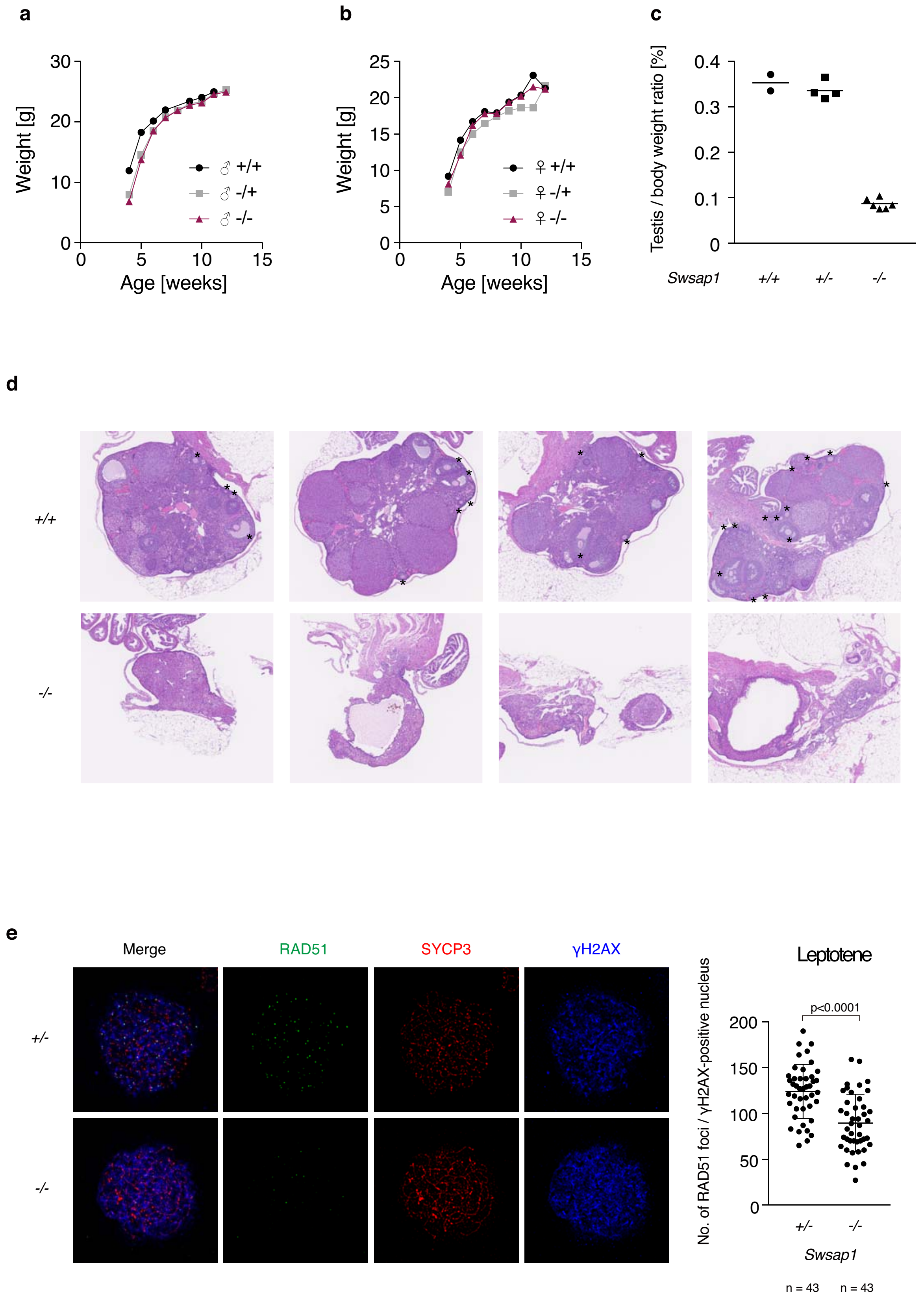


**Supplementary Fig. 4: Meiotic defects and RAD51 assembly defects in *Swsap1*$^{-/-}$ mice**

**a,** Weight analysis of *Swsap1* male mice. Average weights of more than 3 mice for each group are shown. Error bars are not shown to render the graph readable. **b,** Weight analysis of *Swsap1* female mice. Average weights of more than 3 mice for each group are shown. Error bars are not shown to render the graph readable. **c,** Testis weight of wild-type, *Swsap1*$^{+/-}$ and *Swsap1*$^{-/-}$ mice. Testis weight was normalized by body weight. Mean is shown as a bar. **d,** Cross sections of fixed ovary were stained with HE. Several representative images are shown. Asterisks denote developing ovaries. **e,** RAD51, SYCP3 and γH2AX immunofluorescence analysis of leptotene spermatocytes. Left, Representative images of *Swsap1*$^{+/-}$ and *Swsap1*$^{-/-}$ spermatocyte spreads are shown. Right, Quantification of RAD51 foci in γH2AX -positive leptotene spermatocytes. Mean is shown as a bar. Error bars show standard deviations of 43 spermatocytes. Statistics and reproducibility, see accompanying Source data.

**Supplementary Fig. 5**

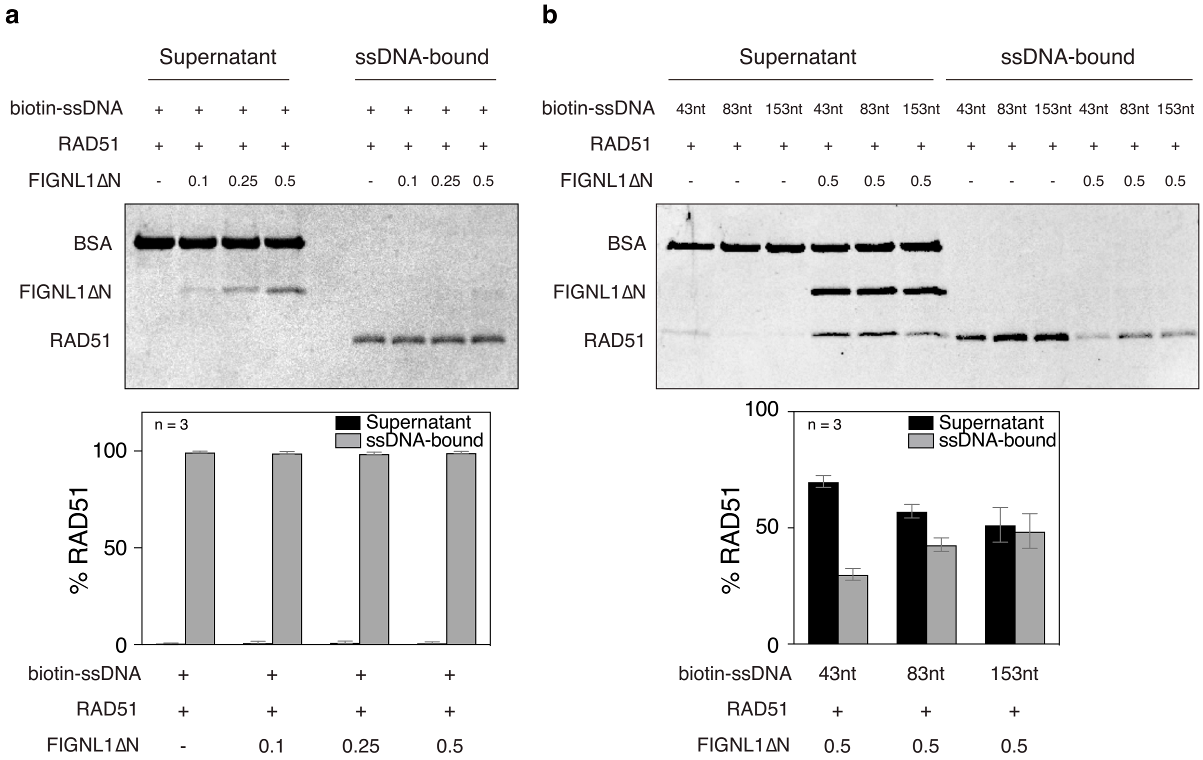


**Supplementary Fig. 5: Property of FIGNL1's RAD51 filament disruption activity.**
**a,** RAD51 disassembly assay in the presence of $Ca^{2+}$. ssDNA pre-bound to RAD51 in the presence of ATP and $Ca^{2+}$ was incubated with an increased concentrations of purified FIGNL1ΔN. After 20 min, supernatants and bound fractions were recovered. Top, a representative SDS-PAGE gel for supernatants and bound fractions stained with CBB. Bottom, Quantification of dissociated RAD51 (Supernatant) and ssDNA-bound RAD51. Intensity of each band of RAD51 was quantified by Imager. The values of RAD51 bands in the supernatant or ssDNA-bound fractions were divided by the total value of RAD51 bands (both in supernatant and bound fractions). Data are mean ± s.d. n=3.; Statistics and reproducibility, see accompanying Source data. **b,** Effect of ssDNA length on RAD51 disassembly. RAD51 filaments formed on 43nt-, 83nt- or 153nt-ssDNA were incubated with 0.5µM FIGNL1ΔN. Top, a representative image of SDS-PAGE gel stained with CBB. Bottom, Quantification of dissociated RAD51 (Supernatant) and ssDNA-bound RAD51. Data are mean ± s.d. n=3.; Statistics and reproducibility, see accompanying Source data.
